# Convergence of unisensory-evoked signals via multiple pathways to the cerebellum

**DOI:** 10.1101/769000

**Authors:** Misa Shimuta, Izumi Sugihara, Taro Ishikawa

## Abstract

The cerebellum receives signals directly from peripheral sensory systems and indirectly from the neocortex. To reveal how these different types of signals are processed in the cerebellar cortex, *in vivo* whole-cell recordings from granule cells and unit recordings from Purkinje cells were performed in mice in which primary somatosensory cortex (S1) could be optogenetically inhibited. Tactile stimulation of the upper lip produced two-phase granule cell responses (with latencies of ∼ 8 ms and 28 ms), for which only the late phase was S1 dependent. Complex spikes and the late phase of simple spikes in Purkinje cells were also S1 dependent. These results indicate that individual granule cells integrate convergent inputs from the periphery and neocortex, and send their outputs to Purkinje cells, which then combine those signals with climbing fiber signals from the neocortex.

## INTRODUCTION

The cerebellum and neocortex are interconnected by a large fiber system (Schmahmann et al., 2019; Watson and Apps, 2019). Although the basal ganglia similarly connect with the neocortex, the cerebellum receives inputs not only from the neocortex but also from the peripheral sensory systems, including tactile, proprioceptive, and vestibular systems (Bostan et al., 2013). How the cerebellum integrates these signals is not well understood.

The cerebellum receives inputs via two types of projection fibers, namely, mossy fibers that project to granule cells and climbing fibers that project to Purkinje cells. A major subgroup of the mossy fibers projects from the pontine nuclei (basilar pontine nuclei and nucleus reticularis tegmenti pontis), which relay signals from the neocortex (Kratochwil et al., 2017; Leergaard and Bjaalie, 2007; Morissette and Bower, 1996). Other mossy fibers originate in the spinal cord and brainstem nuclei, including the trigeminal nuclei, and relay sensory signals directly from the periphery (Sillitoe et al., 2012). Beginning in the 1970s, it was shown that spinal and trigeminal mossy fibers transmitting somatosensory signals from a body part and the corticopontine mossy fibers transmitting signals from the corresponding somatotopic area in the somatosensory cortex terminate in the same areas in the cerebellum (Allen and Tsukahara, 1974; Allen et al., 1974; Bower et al., 1981; Morissette and Bower, 1996; Provini et al., 1968; Tahon et al., 2011). However, at the single-cell level, it is still not clear whether these inputs project to different groups of granule cells (Allen et al., 1974; Tahon et al., 2011) or converge on the same individual granule cells. This issue is important in order to understand the basis of cerebellar computation.

Cerebellar granule cells are small electrically compact cells that receive synaptic inputs from a small number (on average, four) of mossy fibers. Conversely, a single mossy fiber projects to a much larger number (several hundred or more) of granule cells. Thus, Marr and Albus independently proposed similar ideas that the mossy fiber-granule cell system expands the neural representation of information (Albus, 1971; Marr, 1969). This idea, called the expansion recoding hypothesis, assumes that each granule cell receives inputs from a near-random combination of mossy fibers that convey different types of signals, thereby creating numerous combination patterns. Although experimental approaches to test this hypothesis have been hampered by difficulties in tracing individual mossy fibers and recording from single granule cells *in vivo*, more recent studies utilizing neuronal labeling (Huang et al., 2013), *in vivo* electrophysiology (Ishikawa et al., 2015), and *in vitro* electrophysiology combined with neuronal labeling (Chabrol et al., 2015) have demonstrated that mossy fiber inputs conveying different sensory or motor signals converge onto single granule cells. However, the fundamental issue of whether signals evoked by a particular sensory stimulus can converge onto single granule cells after traveling via different pathways has not been investigated.

The cerebellar cortex also receives inputs from climbing fibers, which originate solely from the inferior olivary nuclei. These nuclei comprise a large complex of subnuclei whose inputs have not fully been revealed, but they are known to receive indirect inputs from the neocortex and thus may provide another pathway for cerebrocerebellar communication (Allen et al., 1974; Brown and Bower, 2002; Provini et al., 1968; Ros et al., 2009; Sasaki et al., 1977). However, it has not been demonstrated that the neocortex mediates sensory-evoked climbing fiber responses.

In the present study involving *in vivo* patch clamping and optogenetic manipulation, we provide unequivocal functional evidence that direct trigeminal signals and indirect signals from the primary somatosensory cortex (S1) converge onto the same granule cells. We also show that this integration affects spike outputs of not only the granule cells but also the Purkinje cells. Furthermore, we show that the climbing fiber inputs to Purkinje cells also depend on activity in S1.

## RESULTS

### Photoinhibition of S1 suppresses the late component of cerebellar sensory responses

To investigate how inputs from S1 influence activity in the cerebellar cortex, we recorded field potentials simultaneously from S1 and the granule cell layer (GCL) of the cerebellar cortex in transgenic mice expressing channelrhodopsin 2 (ChR2) in GABAergic neurons (VGAT-ChR2 mice) (Figures 1A and 1B). In mice anesthetized with ketamine/xylazine, a brief (50 ms) air puff applied to the upper lip evoked large responses both in the upper lip area of S1 (3.8–4.5 mm lateral and 0– mm rostral to bregma) and in the GCL of the crus II area. Similarly to that reported in rats (Morissette and Bower, 1996), the responses in the GCL had two peaks, namely, early and late peaks (8.3 ± 0.3 ms and 28.8 ± 0.7 ms [mean ± SEM] from the onset of stimulation, respectively, *n* = 15) that appeared before and after the peak of S1 (25.2 ± 1.2 ms, *n* = 15), respectively (Figures 1C and 1D). When S1 activity was suppressed by focal illumination with blue light in alternating trials, the response of S1 and the late (but not early) component of the GCL response were eliminated (Figures 1C and 1D), indicating that the late component of the GCL response depends on the activity of S1. For this and subsequent experiments, a continuous 150 ms light stimulus was applied during sensory stimulation, but a longer or shorter light stimulus was similarly effective (Figure S1). In similar experiments performed in awake (head-fixed) mice, the early component of the GCL response had two peaks (at 5.1 ± 0.2 ms and 13.0 ± 0.3 ms from the onset of stimulation, *n* = 9), and the S1 response peaked at 14.2 ± 1.0 ms (*n* = 9). The late component of the GCL response, which was measured during a fixed time window (18–23 ms after the onset of stimulation), was eliminated by S1 photoinhibition (Figures 1E and 1F). Of note, the late component was much smaller in awake mice than in anesthetized mice, presumably due to desynchronization in S1 under the awake condition (see Discussion). In a separate set of experiments, anesthetics other than ketamine/xylazine had suppressive effects on both the S1 response and the late component of the GCL response (Figure S2). Therefore, we used ketamine/xylazine in subsequent experiments. Ketamine/xylazine has only minor effects on mossy fiber-granule cell synapses and tonic inhibition of granule cells (Bengtsson and Jörntell, 2007; Duguid et al., 2012).

**Figure 1.**
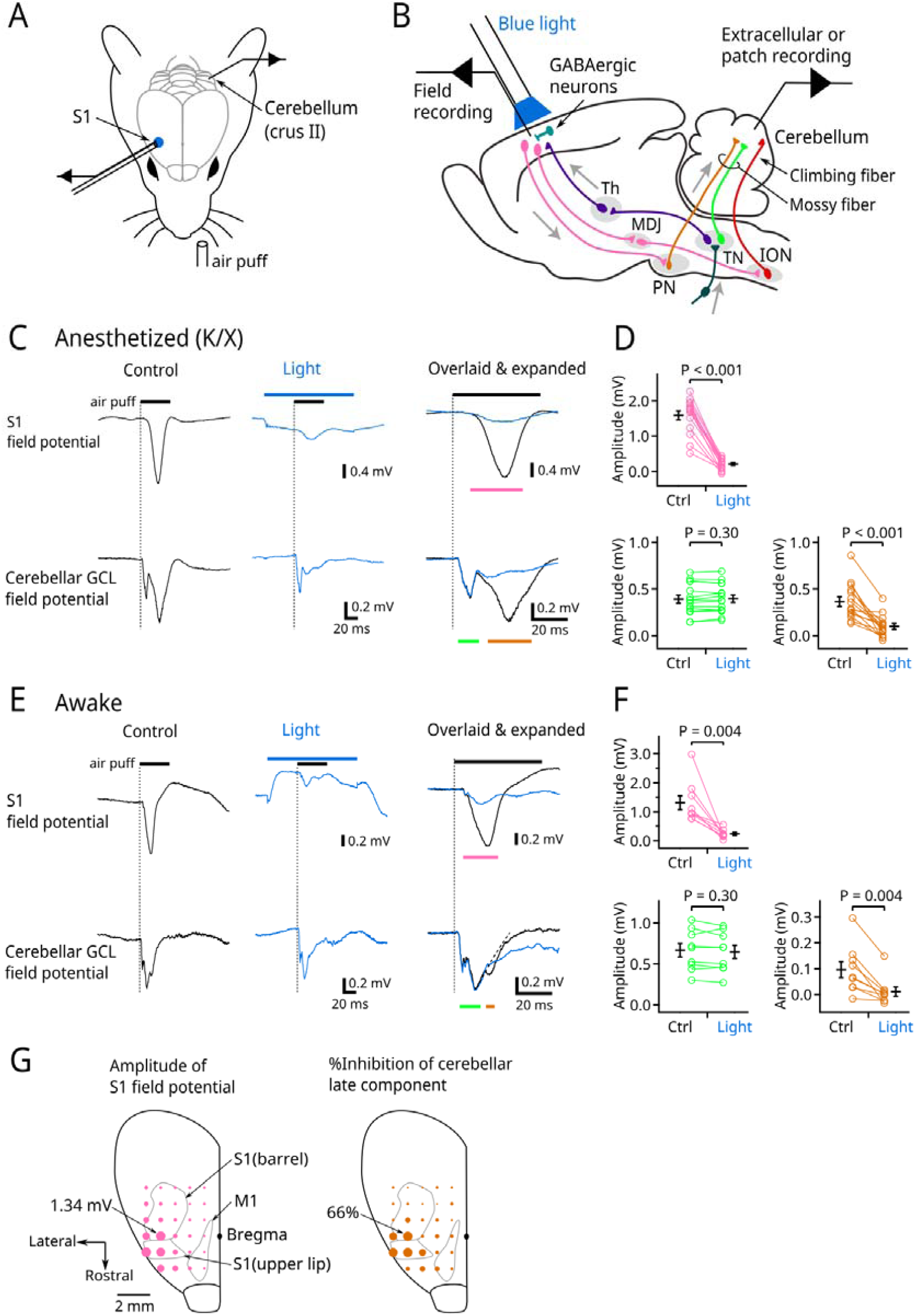
Optogenetic inhibition of S1 suppresses the late component of the cerebellar sensory response. (A) Experimental configuration. Tactile stimulation (air puff) was applied to the left upper lip while field potentials were recorded from the GCL of the ipsilateral crus II area of the cerebellum and the upper lip area of the contralateral S1. S1 was optogenetically suppressed in alternating trials. (B) Schematic of the neural circuit involved in this study. Th, thalamus; MDJ, mesodiencephalic junction; PN, pontine nuclei; TN, trigeminal nuclei; ION, inferior olivary nucleus. (C) Representative recordings from a mouse anesthetized with ketamine/xylazine; 20 traces for each condition were averaged. The same traces are baseline adjusted and overlaid in an expanded time scale in the panels on the right. Black and blue bars indicate the durations of air puff and LED illumination, respectively. The vertical dotted lines indicate the onset of air puff. (D) Pooled data from 15 anesthetized mice. Peak amplitudes were measured during the times indicated by colored bars in (C). (E) Similar to (C) except using an awake mouse; 46 traces for each condition were averaged. The decay time course of the fast component of the cerebellar response was fitted and extrapolated by a single exponential curve (the dashed line) in the panel on the right. (F) Similar to (D) except using nine awake mice. The cerebellar late component was measured after subtracting the decay component of the early response (see Methods). (G) Left, bubble size indicates field potential amplitude at each spot measured in the right cerebral hemisphere. Right, bubble size indicates magnitude of inhibition of the cerebellar late component when blue light was applied to each spot. The cerebellar early component was not affected by photoinhibition at any of these spots. Data from 15 animals were combined. Each spot is the average from 3–15 animals. S1 for upper lip and barrel and the primary motor cortex (M1) are illustrated in accordance with Allen Mouse Brain Atlas and (Mohajerani et al., 2013). Means ± SEMs are presented as black bars and lines, respectively. See also Figures S1 and S2.

To identify the neocortical region involved in the cerebellar response, we performed the same experimental protocol but with the optrode at various loci of the neocortex. The late component of the GCL response was suppressed most effectively by photoinhibition of the locus where the S1 response was largest (3.8 mm lateral and 0 mm rostrocaudal from bregma) but not suppressed by photoinhibition at distant loci, including the primary motor cortex (M1) (Figure 1G). This result suggested that S1 projects directly to the pons, without a relay through other neocortical areas.

### *In vivo* whole-cell recording from granule cells revealed convergent synaptic inputs

The above-described results were consistent with the established theory that at least two distinct groups of mossy fibers project to the GCL of crus II: one directly from the trigeminal nuclei and the other via the cerebropontine pathway (Morissette and Bower, 1996; Steindler, 1985; Woolston et al., 1981). To determine whether these two inputs converge onto individual granule cells, we performed whole-cell patch-clamp recordings in anesthetized mice. In 50% (12/24) of granule cells, excitatory postsynaptic currents (EPSCs) evoked by stimulation of the upper lip had two components (as in the representative cell in Figure 2A), indicating that two types of mossy fibers converge on some individual cells. However, other granule cells had conspicuous EPSCs with only early (12.5%) or late (25%) timing, and the remainder (12.5%) had no response (cutoff, 0.5 events/trial) (Figure 2B). The numbers of EPSC events for the two components did not correlate (*r* = 0.028, *n* = 24), suggesting that these connections were established independently. In these recordings, it was noted that each component often had multiple EPSC events with very short intervals (around 3 ms on average; Figure 2C). This is in line with previous studies showing that a single mossy fiber can fire high-frequency bursts of action potentials that trigger high-frequency EPSCs in granule cells (Chadderton et al., 2004; Rancz et al., 2007). Indeed, in our occasional whole-cell recordings from putative mossy fiber boutons (*n* = 3), action potentials occurred in high-frequency bursts with either early or late timing (Figure S3). Thus, the multiple EPSCs in each component were likely derived from a single mossy fiber. Furthermore, as the interevent intervals for the early and the late EPSC components did not differ (Figure 2C), it is likely that trigeminal and pontine mossy fibers can fire similar high-frequency bursts. However, the fluctuation of the timings (jitter) of the first event was larger in the late component than in the early component (Figure 2D), reflecting a longer multistep pathway for transmission of the late response. However, the amplitude of individual EPSC events was larger for the early components (Figure 2E), suggesting that the synaptic properties (i.e., quantal content and/or quantal size) may differ between these two types of synaptic inputs, as reported previously for the various vestibular inputs (Chabrol et al., 2015). As expected, the late component of EPSCs was mostly eliminated by optogenetic suppression of S1 (Figure 2F), confirming that the late component signal is derived from S1. Similar results were obtained when synaptic charge (the area over the curve), instead of event number, was measured as an indicator of synaptic strength (Figures S4A and S4B). Furthermore, spontaneous EPSCs were partially blocked by the suppression of S1 (Figure S5). These results indicate that granule cells receive convergent inputs from two types of mossy fibers, although the balance of these inputs varies between cells. Additionally, we recorded inhibitory postsynaptic currents (IPSCs) in a small number of granule cells (*n* = 4) to examine feed-forward inhibition from Golgi cells. IPSCs also exhibited two components, and the late component was suppressed by light illumination of S1 (Figures S4C and S4D).

**Figure 2.**
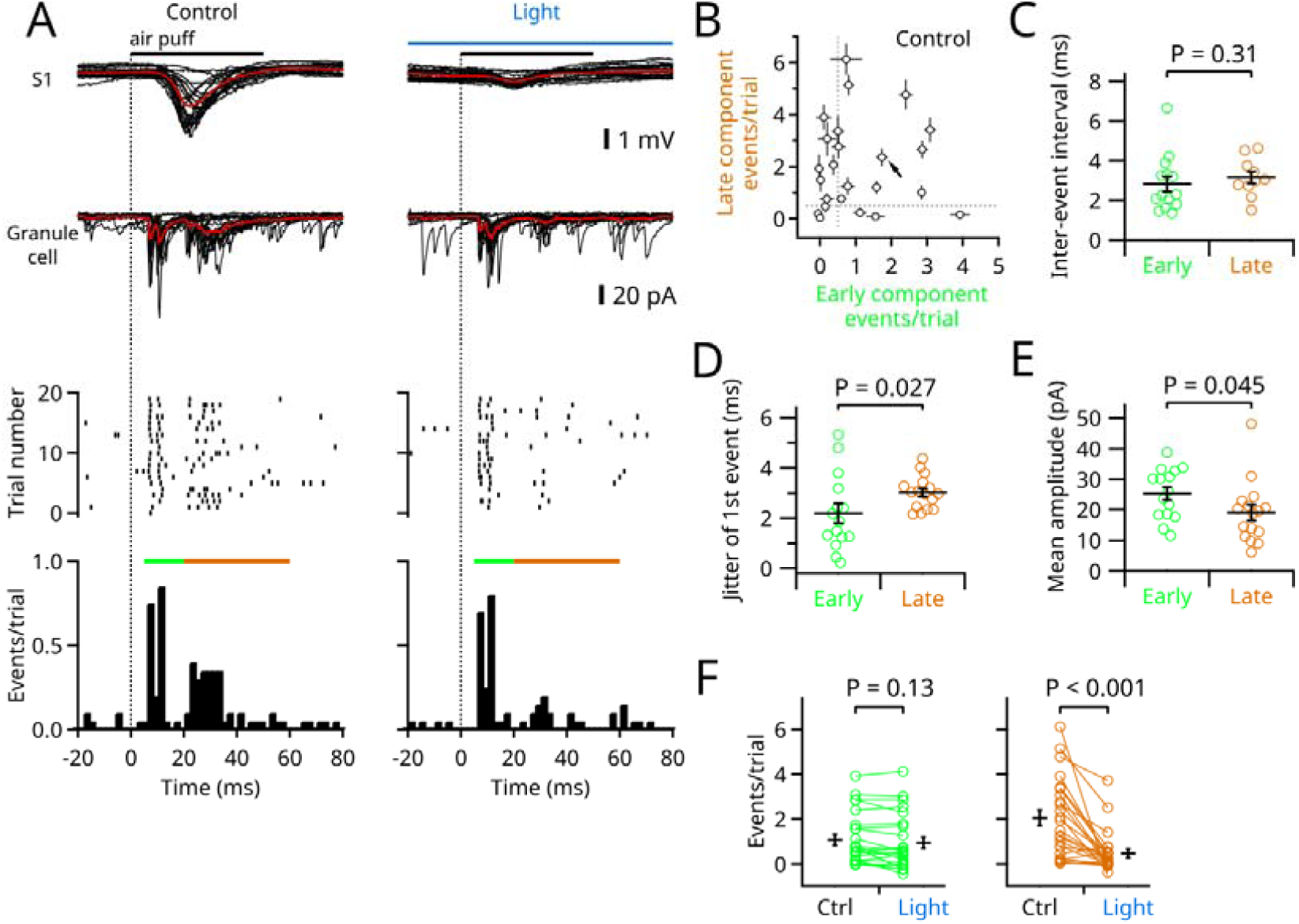
*In vivo* whole-cell recording from granule cells revealed convergent synaptic inputs. (A) Representative recording; simultaneous field potential recording from S1 (top) and whole-cell voltage clamp recording from granule cell (lower); 20 consecutive traces are overlaid. The average traces are in red. Detected EPSC events are shown in raster plots (middle) and time histograms (bottom). Green and brown bars indicate the early and the late phases. Trials under control (left) and the light (right) conditions were interleaved. (B) The numbers of evoked EPSC events in the early and the late phases were compared in 24 cells. The numbers were corrected for baseline spontaneous events by subtraction in this and the following panels. For qualitative description, event numbers of <0.5 events/trial (dotted lines) were defined as no response. Error bars indicate SEMs (trial-by-trial fluctuation). The arrow points to the cell shown in (A). (C) The median interevent interval of each cell was compared (*n* = 14 in the early and *n* = 10 in the late). (D) Trial-by-trail fluctuations (measured as standard deviation) of the timing of the first event during each time period were compared (*n* = 15 in the early and *n* = 16 in the late). (E) The mean amplitudes of EPSCs in the early and late phases were compared (*n* = 15 in the early and *n* = 16 in the late). (F) EPSC event numbers were compared for the early phase (left) and the late phase (right) (*n* = 24). Means ± SEMs are presented as black bars and lines, respectively. See also Figures S3, S4, and S5.

### Spike output of single granule cells *in vivo*

To investigate how synaptic inputs trigger action potentials in granule cells, we recorded membrane potentials in current clamp (Figure 3A). The resting potentials of granule cells varied widely (from −96.0 to −50.8 mV; mean, −72.9 ± 3.4 mV, *n* = 16). In cells that had a relatively hyperpolarized resting potential, the excitatory postsynaptic potential (EPSP) did not reach the spike threshold (Figure 3B), indicating that a single tactile stimulus generates a spike in only a subset of the granule cells. As the resting potentials of granule cells are modulated by multiple factors, such as tonic inhibition from Golgi cells and potassium channel function (Duguid et al., 2012; Hamann et al., 2002; Millar et al., 2000), we applied steady depolarizing current (up to 20 pA) to 9 of the 16 cells, thereby raising the average resting potential to −57.2 ± 1.9 mV (*n* = 16, including seven cells without adjustment) and increasing their propensity to fire in response to tactile stimulation of the upper lip (11/16 cells fired action potentials; cutoff, 0.5 spikes/trial) (Figures 3A and 3B). In this condition, the granule cells fired during the early or late phase (Figure 3C). As expected, action potentials in the late phase were eliminated by photoinhibition of S1 (Figure 3F). Investigation of the relationship between synaptic inputs and firing outputs showed that the number of action potentials evoked in the early phase correlated with the number of EPSC events during the same period (*r* = 0.77, *n* = 13) (Figure 3D, left). However, the correlation was less clear in the late phase (*r* = 0.53, *n* = 13) (Figure 3D, right), suggesting that factors other than instantaneous synaptic inputs may be involved. Indeed, depolarization during the early phase was maintained through the late phase (Figure 3E). This suggests that action potential firing in the late phase may be facilitated by temporal summation. A simulation in a realistic computational model of a granule cell (Diwakar et al., 2009) showed such a facilitation (Figure S6).

**Figure 3.**
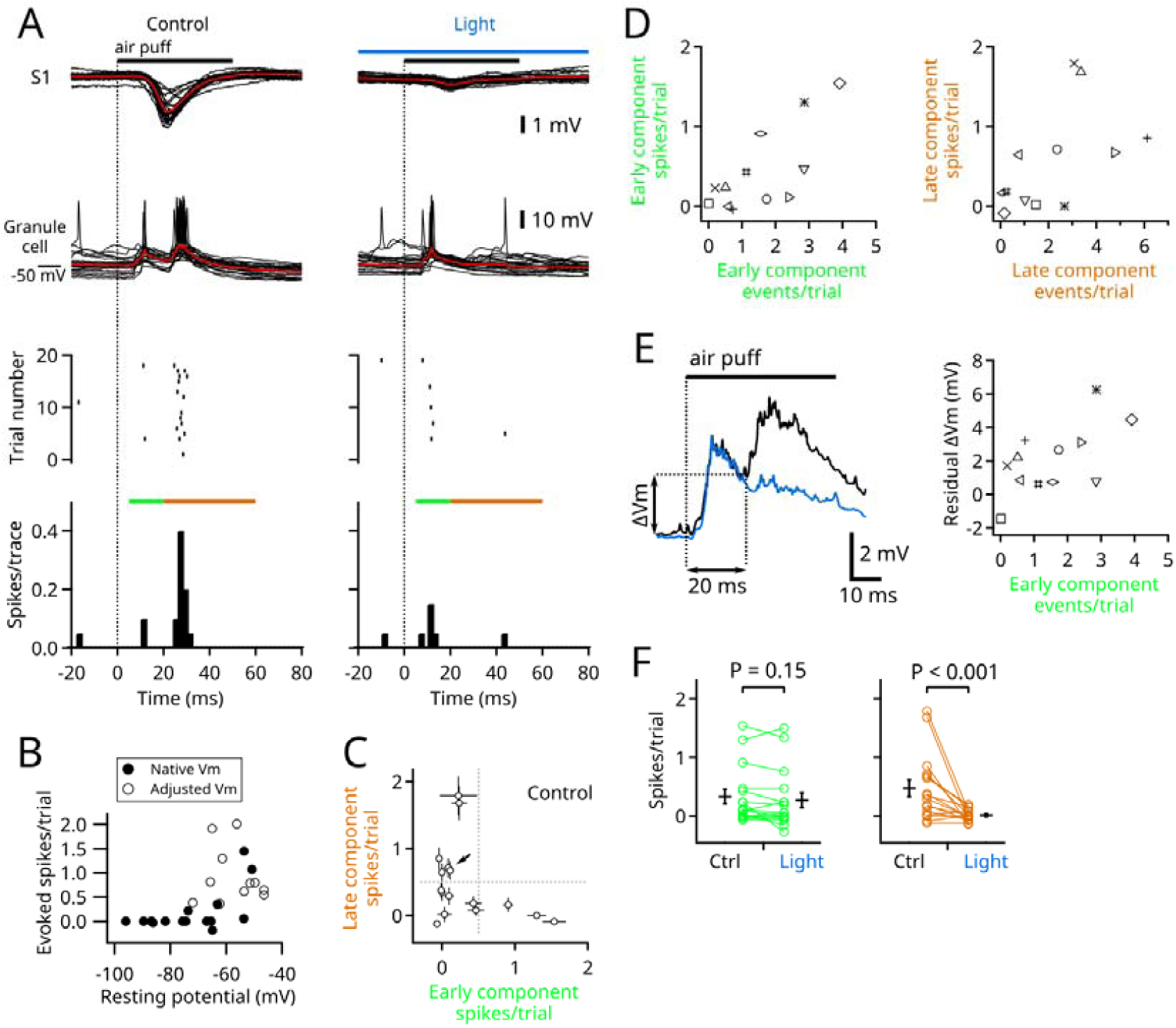
Spike output of single granule cells *in vivo*. (A) Representative recordings of the same cell as in Figure 2A; simultaneous field potential recordings from S1 (top) and whole-cell current clamp recordings from a granule cell (lower); 20 traces are overlaid. The average traces are in red. Detected spikes are shown in raster plots (middle) and time histograms (bottom). Green and brown bars indicate the early and late phases. The trials under control (left) and light (right) conditions were interleaved. (B) Numbers of evoked spikes (sums of the early and the late responses) plotted against the native resting potentials (filled circles, *n* = 16). In a subset of cells that had hyperpolarized native resting potentials (<65 mV), a steady depolarizing current was applied to facilitate firing (open circles, *n* = 9). The spike numbers were corrected for baseline spontaneous firing by subtraction in this and the following panels. (C) The numbers of evoked spikes in the early and the late phases were compared in 16 cells, including three cells that lacked voltage clamp recording. For qualitative description, event numbers of <0.5 spikes/trial (dotted lines) were defined as no spikes. Error bars indicate SEMs (trial-by-trial fluctuation). The arrow points to the cell shown in (A). (D) Numbers of evoked spikes in the early phase (left) and late phase (right) were plotted against the numbers of EPSCs during the same time periods. Individual cells (*n* = 13) are plotted as different marks common in (D) and (E). (E) Left, averaged traces of all 16 granule cells in current clamp mode. Black, control; blue, photoinhibition of S1. Depolarization at 20 ms after the onset of stimulation was measured as residual depolarization. Right, the residual depolarization of each cell is plotted against the number of EPSCs in the early phase. (F) Evoked spike numbers were compared for the early phase (left) and the late phase (right) (*n* = 16). Means ± SEMs are presented as black bars and lines, respectively. See also Figure S6.

### Sensory-evoked responses in granule cells in various parasagittal bands

The cerebellar recording electrodes in the above-described experiments were positioned coarsely around the 6+ band in crus II. To characterize the distribution of trigeminal and corticopontine projections in crus II, we used Aldoc-Venus mice in which the various bands can be visualized (Figure 4B). Field potentials were recorded from S1 simultaneously with *in vivo* whole-cell voltage clamp recordings from the cerebellar granule cells in 5+, 5-, 6+, and 7+ bands. As observed in VGAT-ChR2 mice, some granule cells had EPSCs in both early and late time periods, whereas others had EPSCs in only one of these time periods (Figure 4A). Although early and late responses were detected in all bands, the 7+ band had the fewest early component EPSC events, and the fewest late component events were recorded in the 5-band (Figures 4C and S7B). Consistent results were obtained with field potential recordings (Figure S7A). These results indicate that the putative trigeminal and corticopontine mossy fiber inputs are variably distributed over multiple bands in crus II. Furthermore, trial-by-trial analysis revealed a correlation between the amplitude of the S1 response and the number of EPSC events in the late component but not in the early component (Figures 4D and 4E), confirming that only the late component is S1 dependent. Similar to that observed in VGAT-ChR2 mice, the mean EPSC amplitude was larger and the jitter of the first event was smaller in the early response than in the late response, and the interevent intervals did not differ (Figures S7C–E).

**Figure 4.**
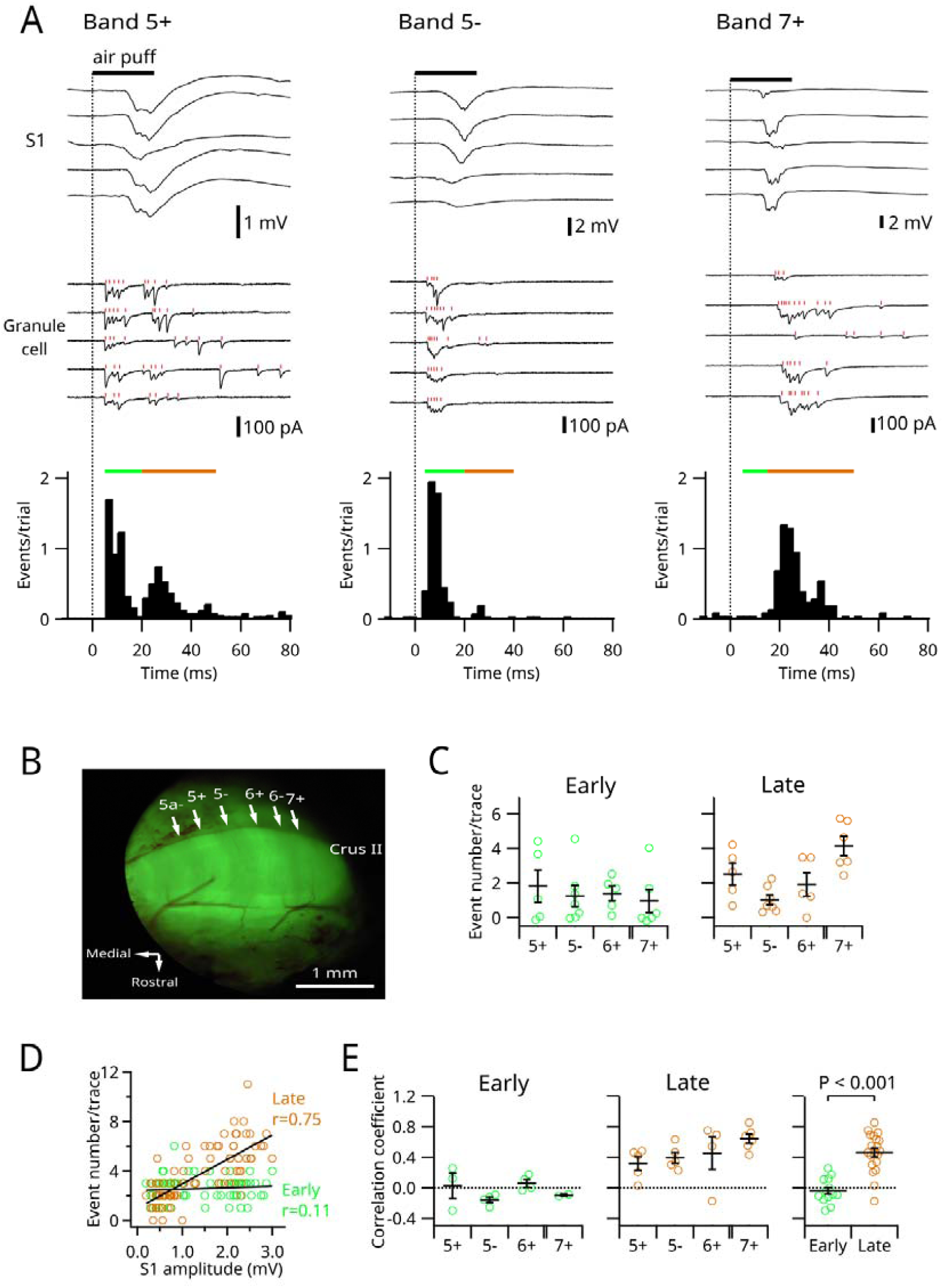
Sensory-evoked responses in granule cells in various parasagittal bands. (A) Representative recordings; simultaneous field potential recordings from S1 (top) and whole-cell voltage clamp recordings from a granule cell (lower); five traces are displayed. Overlaid raster plots in red indicate the timing of detected EPSC events. Time histograms are shown in the bottom panels. Green and brown bars indicate the early and the late phases. (B) Fluorescent macroscopic image of the cerebellar crus II area after removal of the dura mater. (C) The numbers of EPSC events in different bands in the early (left) and the late (right) phases. (D) Data from a sample cell in the 6+ band. The EPSC event numbers in the early (green) and late (brown) phases in every trial were plotted against the amplitudes of S1 field responses. Correlation coefficients were calculated for each phase. (E) Correlation coefficients of all cells are plotted. The right panel shows pooled data from all bands. Means ± SEMs are presented as black bars and lines, respectively. See also Figure S7.

### Effect of S1 photoinhibition on SS and CS in Purkinje cells

We next investigated how the suppression of S1 affects the firing of Purkinje cells, which are downstream of granule cells in the cerebellar circuit (Bower and Woolston, 1983). In general, simple spikes (SS) in Purkinje cells are generated spontaneously but are affected by excitatory synaptic inputs from parallel fibers (i.e., granule cell axons) and inhibitory synaptic inputs from molecular layer interneurons, whereas complex spikes (CS) are triggered solely by climbing fiber inputs. Under control conditions (without S1 inhibition), there were four phases of SS in response to stimulation of the upper lips of VGAT-ChR2 mice, namely, early excitatory (within 10 ms after stimulation onset), early inhibitory (around 10–20 ms), late excitatory (around 20–30 ms), and late inhibitory (30-60 ms) phases (Figures 5A–5D). CS coincided with the late inhibitory phase of SS.

**Figure 5.**
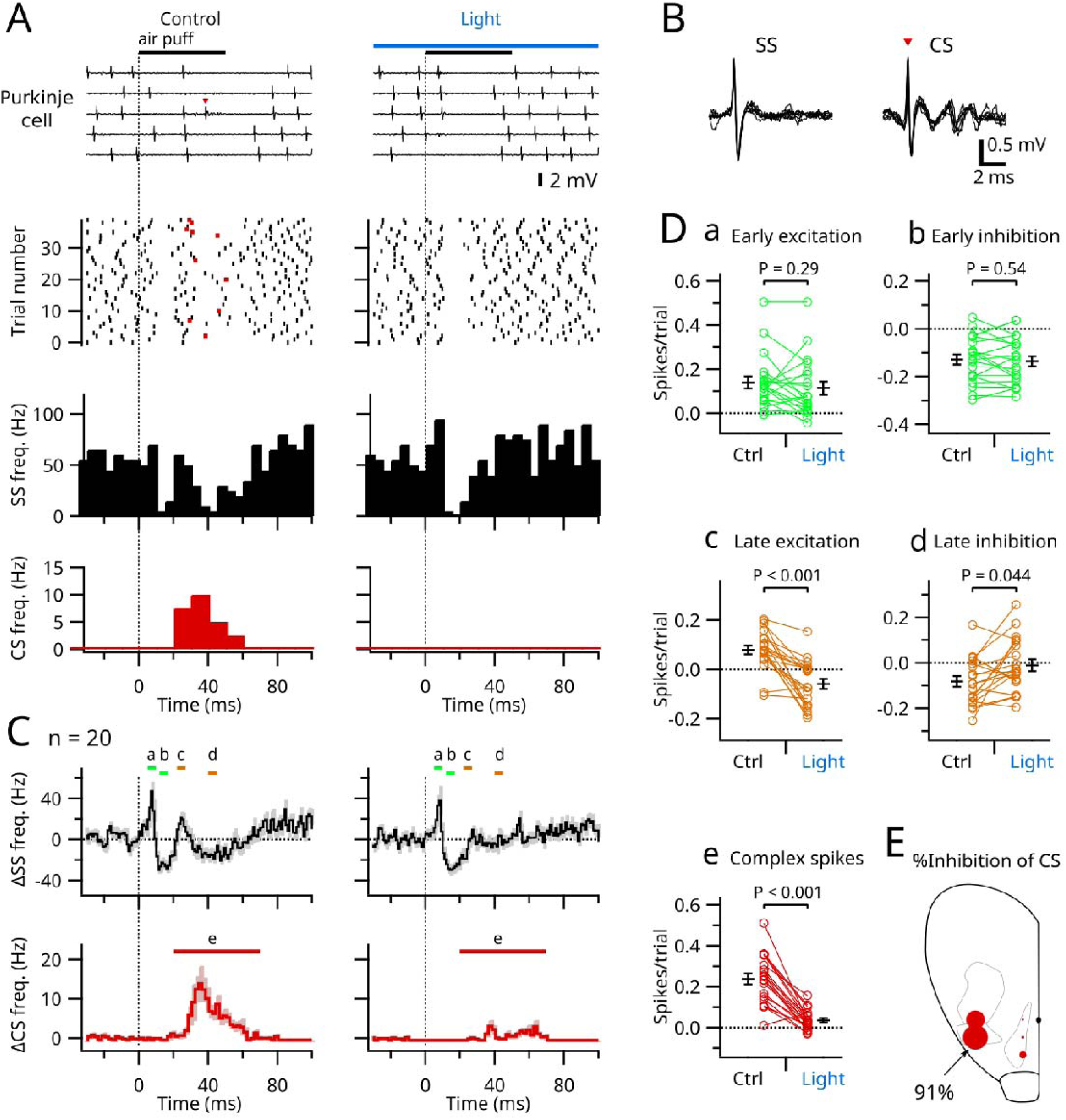
Effects of S1 photoinhibition on SS and CS in Purkinje cells. (A) Representative extracellular unit recordings of a Purkinje cell; five traces are displayed (top). Simultaneous field potential recordings from S1 are not shown. SS (black) and CS (red) are plotted in the raster plots (lower). Separate time histograms of SS (middle) and CS (bottom) are shown. (B) Enlarged views of SS and CS of the unit shown in (A); six traces in each were aligned and overlaid. (C) Averaged time histograms of SS (1 ms bin) and CS (2 ms bin) from 20 Purkinje cells. Shading indicates SEMs. Vertical dotted lines indicate the onset of stimulation as in (A). (D) Spike numbers were measured during the times (a–e) indicated by colored bars in (C). Means ± SEMs are presented as black bars and lines, respectively. (E) Bubble size indicates magnitude of inhibition of the cerebellar CS when the blue light was applied to each spot in the right cerebral hemisphere. Only five spots were tested. Data from six animals were combined. Each spot is the average from 3–6 animals. See also Figures S8 and S9.

Photoinhibition of S1 did not affect the early phases of SS but eliminated the late excitatory and inhibitory phases (Figures 5A–5D). Interestingly, the CS response was also abolished. These results suggest that the direct trigeminal inputs to granule cells trigger the early excitatory and inhibitory phases of SS, the latter of which presumably results from the activation of molecular layer interneurons (Blot et al., 2016; Jelitai et al., 2016). These results also suggest that the late excitatory responses of SS reflect inputs from S1. However, the late inhibitory response could reflect either an interneuronal effect or a pause after CS. As CS did not always occur under the control condition, we were able to separate traces with CS from those without (Figure S8). The late inhibition phase of SS was larger in traces with CS than in those without CS, suggesting that the late inhibition largely reflects the pause after CS. We also tested the effects of photoinhibition at multiple loci in the neocortex and found that suppression of the upper lip area of the contralateral S1 was most effective in blocking CS in Purkinje cells (Figure 5E). Furthermore, suppression of S1 with a long (10 s) light stimulus inhibited spontaneous CS firing and triggered rebound activation after the light was turned off, suggesting that the spontaneous activity of the inferior olive is under strong control of S1. However, this photoinhibition had no effect on SS firing (Figure S9), suggesting that S1 activity is not directly linked to the spontaneous activity of granule cells (Figure S5) in line with a previous report (Ros et al., 2009).

## DISCUSSION

By taking advantage of the high temporal resolution of electrophysiology and optogenetics, we obtained functional evidence that cerebellar granule cells in crus II receive inputs directly from the periphery and indirectly via S1. We found that approximately half of the granule cells receive convergent signals from both pathways. We also showed that the olivocerebellar inputs to these Purkinje cells come through the same area in S1.

### Optogenetic analysis of the neural pathways

We adopted an optogenetic method in which photostimulation of GABAergic neurons expressing ChR2 can suppress the activity of specific areas of the neocortex (Guo et al., 2014; Li et al., 2015, 2016). This method is based on the fact that most GABAergic neurons in the neocortex are local interneurons. Although 0.5% of GABAergic neurons project to other cortical and subcortical areas (Tamamaki and Tomioka, 2010), it is unlikely that such long-range GABA projections contributed to our findings because light illumination was effective only in specific areas in the neocortex (Figures 1G and 5E). Moreover, plenty of anatomical and physiological evidence indicates that the basal pons relays signals from S1 to the cerebellar cortex (Leergaard and Bjaalie, 2007; Leergaard et al., 2006; Morissette and Bower, 1996; Watson and Apps, 2019; Welker, 1987). Therefore, it is highly likely that the late component of the cerebellar response in our study was transmitted via the cortico-ponto-cerebellar pathway.

### Convergence of mossy fiber inputs to granule cells

In contrast to the conventional view of Marr and Albus (Albus, 1971; Marr, 1969) that granule cells receive inputs from mossy fibers carrying different types of information, it was recently proposed that individual granule cells receive only one type of input (Bengtsson and Jörntell, 2009; Dean et al., 2010; Spanne and Jörntell, 2015). Although both trigeminal and corticopontine signals were evoked by the same tactile stimulation in our study, these signals are fundamentally different because they can be differentially modulated on route to the cerebellum. For instance, neocortical states (such as anesthesia and wakefulness) affected them differently (Figure 1), and they exhibited different trial-to-trial fluctuations (Figure 4). Furthermore, as the S1 in mice sends efferent motor signals directly to the brain stem (Matyas et al., 2010), the corticopontine signals we observed may in fact be efferent copies of a motor command. Thus, since the distinctions between trigeminal and corticopontine signals are substantial, our results are in line with the conventional idea proposed by Marr and Albus.

As granule cells received either or both types of inputs, it is difficult to draw a unified view for these connections. For the subpopulation of granule cells that receive both, our results (Figure 3E) indicate that depolarization during the early phase may facilitate firing in the late phase. In such cases, the granule cells represent coincident detectors (i.e., an “AND” gate) and pattern recoders as proposed by Marr and Albus. By contrast, granule cells that fire in response to a burst of synaptic input from either a direct or S1-mediated connection may work as a simple relay or a frequency filter (Bengtsson and Jörntell, 2009; Dean et al., 2010; Rancz et al., 2007). We speculate that a granule cell can become an “OR” gate if both direct and S1-mediated inputs are strong enough to trigger action potentials independently. An experimental system in which trigeminal and corticopontine pathways can be activated independently is required to test this.

Previous morphological studies showed that aldolase-c-positive bands have more corticopontine inputs than trigeminal or spinal ones, and that aldolase-c-negative bands have the opposite trend (Biswas et al., 2019; Na et al., 2019), although there may be some overlap (Valera et al., 2016). Our present results (Figures 4 and S7) are consistent with those studies, confirming that different types of mossy fibers tend to innervate different compartments at the macroscopic level. However, at the single-cell level, the strengths of these two types of inputs were neither positively nor negatively correlated (Figure 2B), suggesting that two independent mossy fibers synapse with a granule cell by chance, without attracting or repelling each other (Solowska et al., 2002). The randomness of these connections is consistent with the expansion recoding hypothesis of granule cells proposed by Marr and Albus (Cayco-Gajic and Silver, 2019).

An electrophysiological study using extracellular recording suggested that the GCL has a stratified organization in which the trigeminal mossy fibers are located deeper than the corticopontine mossy fibers (Tahon et al., 2011). Although our results do not directly contradict this idea, they strongly suggest that the trigeminal and corticopontine mossy fibers largely overlap in the GCL even if they tend to be distributed at different depths. Future studies, such as those using functional cellular imaging (Giovannucci et al., 2017; Wagner et al., 2017, 2019), may elucidate the detailed organization of the GCL.

As there are extensive studies regarding inhibitory inputs to granule cells from Golgi cells (Duguid et al., 2012, 2015; Tabuchi et al., 2018; Vos et al., 1999), we performed only a few experiments on IPSCs in this study. We found that, like EPSCs, the IPSC events had two phases. These currents were delayed in relation to EPSCs by only a few milliseconds, reflecting rapid feed-forward inhibition from Golgi cells. The importance of the excitation/inhibition balance in granule cells has been discussed elsewhere (D’Angelo et al., 2013; Mapelli et al., 2014).

### Convergence of inputs to Purkinje cells

We found that Purkinje cells receive signals from the neocortex not only via the mossy fiber-parallel fiber pathway but also via climbing fibers. This finding is consistent with anatomical observations that Purkinje cells in the 6+ compartment (D1 band) receive inputs from climbing fibers from the ventral principal olive, which receives inputs from the area parafascicularis prerubralis in the mesodiencephalic junction, which, in turn, receives projections from the neocortex (Carlton et al., 1982; Naus et al., 1985; Sugihara and Shinoda, 2004). Our present study is also in line with classical physiological studies showing a convergence of mossy and climbing fiber signals originating from the same neocortical areas in cats and monkeys (Allen et al., 1974; Provini et al., 1968; Sasaki et al., 1977) as well as a more recent study showing that injections of lidocaine into the somatosensory cortex delayed the timing of CS in crus II in rats (Brown and Bower, 2002). Still, our present study is the first to clearly demonstrate that S1 mediates sensory-evoked climbing fiber responses. In contrast to what we observed, a recent study (Kubo et al., 2018) reported that inhibition of the somatosensory cortex in mice did not inhibit CS. The reason for this discrepancy is unclear but may involve minor differences in sensory stimulation, optogenetic methods, or the sites recorded.

The physiological significance of this cortio-olivo-cerebellar connection is intriguing, especially given the parallel cortico-ponto-cerebellar connection. This circuitry suggests that the neocortex exerts powerful control over cerebellar signal processing. For instance, it may be possible that signals from the neocortex induce synaptic plasticity at synapses on Purkinje cells when parallel and climbing fibers are activated simultaneously via the respective cortico-ponto-cerebellar and cortio-olivo-cerebellar pathways. Such top-down signaling may occur not only during wakefulness but also during sleep as offline activity (Miyamoto et al., 2016; Ramanathan et al., 2015).

### Limitations and future directions

One limitation of the present study is that the data were largely obtained from animals anesthetized with ketamine/xylazine, which has a suppressive effect on parallel fiber-Purkinje cell synapses (Bengtsson and Jörntell, 2007; Duguid et al., 2012). However, the sensory stimulation applied was able to evoke SS in Purkinje cells. Ketamine/xylazine also synchronizes activity in the neocortex (Pachitariu et al., 2015), which may enhance the size and distribution of cortical responses to a punctuate stimulus (Harris and Thiele, 2011; Poulet and Crochet, 2018). The fact that S1-mediated signals observed in the cerebellum were clearer (larger and better isolated) in anesthetized animals than in awake animals (Figure 1) may be attributable to this effect. As the precise mechanisms of action of the anesthetics are not known, further studies are required to understand how brain states, including natural sleep, affect cerebrocerebellar communication.

A second limitation is that we did not have a method to specifically block the trigemino-cerebellar pathway. A future challenge is to manipulate the trigeminal neurons projecting to the cerebellum without affecting those projecting to the thalamus. This may be possible in principle as those neurons may be different populations in the trigeminal nuclei (Bennett-Clarke et al., 1992; Zhang et al., 2018). On a related note, we do not know whether the early phase inputs from the trigeminal nuclei to the cerebellar granule cells were monosynaptic or polysynaptic. Although these responses had very short latencies, it is possible that rapid polysynaptic inputs, relayed within the trigeminal nuclei or in other brainstem areas, contributed. Indeed, the early responses in awake mice had two clear peaks. In this study, we collectively referred to these early responses as the direct trigeminal response to distinguish them from the indirect S1-mediated response.

Finally, in this study, the only sensory stimulus was tactile stimulation (air puff) of the upper lip. We adopted this form of stimulation because it gave large responses in both S1 and the cerebellum. However, tactile stimulation of other body parts or stimulation of other modalities may similarly evoke multipathway responses in the cerebellar cortex. Given the extensive connections between the neocortex and the cerebellum (Leergaard and Bjaalie, 2007; Proville et al., 2014), it is likely that similar circuits exist for the sensation of other types of stimulation.

## METHODS

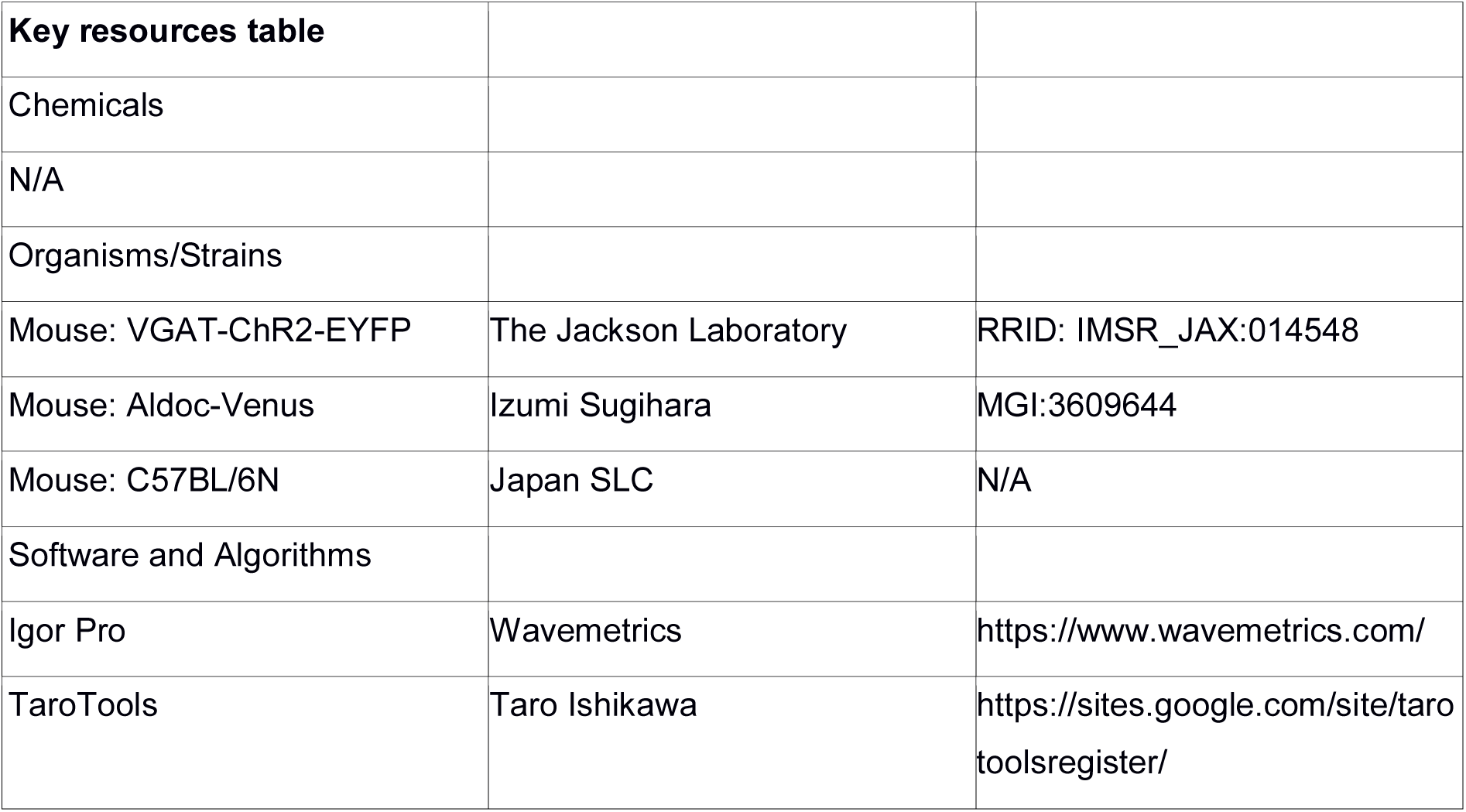

### Animals

All animal procedures were approved by the Institutional Animal Care and Use Committee of The Jikei University (no. 2015-054 and 2017-001) under the Guidelines for the Proper Conduct of Animal Experiments of the Science Council of Japan (2006).

Hemizygous transgenic mice (3–8 weeks old, male and female) expressing modified ChR2 (hChR2-H134R) fused with YFP in GABAergic neurons via the vesicular GABA transporter promoter/enhancer (VGAT-ChR2 mice, stock no. 014548; Jackson Laboratory) (Zhao et al., 2011) were used in photoinhibition experiments. Heterozygous knock-in mice (3–5 weeks old, male and female) expressing Venus fluorescent protein via the aldolase-C promoter (Aldoc-Venus mice, MGI:3609644) (Fujita et al., 2014) were used to visually identify the cerebellar parasagittal zones.

### Surgery

The mice were anesthetized with an initial dose of a mixture of ketamine (86 mg/kg body weight) and xylazine (10 mg/kg), and supplemented with a continuous infusion of ketamine (70 mg/kg/h) and xylazine (8 mg/kg/h) via a syringe pump to maintain a stable level of anesthesia with free breathing. The heads of mice were fixed in position via a head post glued onto the skull. Core body temperature was maintained at around 37□C with an isothermic feedback heating pad. After removing the overlaying skin and muscles, a craniotomy was performed over the left cerebellar crus II area and the right cerebral somatosensory area. After removing the dura, the exposed brain surface was kept moist with a HEPES-buffered saline containing (in mM) NaCl 150, KCl 2.5, HEPES 10, CaCl_2_ 2, and MgCl_2_ 1 (pH adjusted to 7.4 with NaOH). To record from awake mice, the head post was attached during a surgery 3 days before recording. Craniotomies were performed under isoflurane anesthesia on the recording day. The mice were habituated to the recording environment for >1 h before the start of recording. The duration of the recording session was <5 h for each animal.

### Recording

Whole-cell *in vivo* patch-clamp recordings were performed via a resistance-guided (blind) method as previously described (Ishikawa et al., 2015; Rancz et al., 2007). Voltage clamp and current clamp recordings were made from granule cells at a depth of 200–400 μm in crus II of the cerebellar cortex by using a Multiclamp 700B amplifier (Molecular Devices). Data were low-pass filtered at 6 kHz and acquired at 50 kHz using a USB-6259 interface (National Instruments) and Igor Pro with NIDAQ Tools MX S(WaveMetrics). The internal solution contained (in mM) K-methanesulfonate 135, KCl 7, HEPES 10, Mg-ATP 2, Na_2_ATP 2, Na_2_GTP 0.5, and EGTA 0.05 or 0.1 (pH adjusted to 7.2 with KOH), giving an estimated chloride reversal potential of −69 mV. This enabled excitatory synaptic currents to be observed in isolation by voltage clamping at −70 mV and inhibitory synaptic currents to be observed at 0 mV. The liquid junction potential was not corrected. In a subset of experiments, biocytin 0.5% was added to the internal solution. The granule cells were identified by a small membrane capacitance (C_m_) (less than 7 pF) and a lack of periodic spontaneous firings. The mossy fiber boutons were identified by the occurrence of high-frequency bursts and a lack of synaptic potentials (Rancz et al., 2007). Patch pipettes had resistances of 5–8 MΩ, and series resistances (R_s_) were typically 30–50 MΩ. Cells with high R_s_ were excluded to keep the access time constant (R_s_·C_m_) at <0.3 ms. Extracellular unit recordings from Purkinje cells were made with glass electrodes (5 MΩ) filled with saline. The Purkinje cells were identified by the occurrence of SS and CS. Field potential recordings were conducted using a tungsten electrode (1 MΩ, TM31A10; WPI). Neocortical field potentials were recorded at a depth of 550–650 μm in the primary somatosensory area for the upper lip (3.8 mm lateral and 0 mm rostrocaudal from bregma) unless otherwise noted. Cerebellar field potentials were recorded in the GCL at a depth of 300–500 μm in the crus II area. In VGAT-ChR2 mice, the somatosensory cortex was optogenetically inactivated by illuminating the surface with an optic fiber (0.39 NA, Ø400 μm, FT400EMT; Thorlabs) coupled to a high-power blue LED (470 nm, M470D2; Thorlabs). The optic fiber was bundled with a single tungsten electrode, and the tip of the electrode was 800 μm ahead of that of the optic fiber so that the tip of the optic fiber was above the surface of the brain during the recording. To record from visually identified bands in Aldoc-Venus mice, the angle of electrode was carefully adjusted to be perpendicular to the brain surface, and the center of each band was targeted. However, due to the limited spatial specificity inherent in field potential recording (Kajikawa and Schroeder, 2011), contamination of field potentials from neighboring bands was not fully excluded. For whole-cell recordings from those mice, only 5+, 5-, 6+, and 7+ bands, which were relatively wide and easy to identify, were used. Tactile stimulation (air puff, 50 ms, 90 mmHg at the source) was applied to the left upper lip with a solenoid valve (FAB31-8-3-12C-1; CKD, Aichi, Japan) controlled via the USB-6259 interface as described above.

### Analysis

To analyze field potential recordings, the peak amplitude from the baseline was measured. In ketamine/xylazine anesthetized mice, the baseline for the cerebellar late component was set at the beginning of the late component, which could be readily identified. In awake mice and mice anesthetized with a mixture of medetomidine (0.3 mg/kg), midazolam (4 mg/kg), and butorphanol (5 mg/kg) or with isoflurane, the late component was measured as the mean amplitude 18–23 ms from the stimulation onset after subtracting the extrapolated decay component of the early response (a single exponential curve common under control and photoinhibited conditions). EPSCs and action potentials in whole-cell recordings were detected using TaroTools (https://sites.google.com/site/tarotoolsregister/), a threshold-based algorithm in Igor Pro (WaveMetrics). The event number (for both synaptic events and action potentials) evoked by stimulation was counted (baseline subtracted) in a time window adjusted for each cell to include all evoked events but to minimize contamination of spontaneous events. The beginning of the time window was adjusted to between 0 and 5 ms from the onset of stimulation, and the end of the time window was between 50 and 60 ms. The border between the early and the late phases was adjusted to between 16 and 20 ms so that the border matched the valley of the time histogram. The synaptic charge was measured as the integral of the averaged current trace (baseline subtracted and sign reversed). The latency of the sensory response was defined as the time from the stimulus onset to the first EPSC event. Data are presented as means ± SEMs. Statistical significance was tested by using Wilcoxon signed-rank tests for paired samples and Wilcoxon– Mann–Whitney tests for independent samples.

## Acknowledgments

The authors are grateful to Toshihiko Momiyama, Soichi Nagao, and Hirofumi Tokuoka for their generous support, to Tadaharu Tsumoto and Kazuhiro Sohya for their help in obtaining VGAT-ChR2 mice, and to Miwa Takagi for technical assistance. This work was supported by JSPS KAKENHI (grant number 18K06529), MEXT-Supported Program for the Strategic Research Foundation at Private Universities (S1311009), and the Jikei University Research Fund.

## Author contributions

Conceptualization, M.S. and T.I.; Methodology, M.S. and T.I.; Software, T.I.; Formal Analysis, T.I.; Investigation, M.S. and T.I.; Resources, I.S.; Writing –Original Draft, T.I.; Writing –Review & Editing, M.S. and I.S.; Supervision, I.S. and T.I.; Project Administration, T.I.; Funding Acquisition, M.S., I.S. and T.I.

## Declaration of interests

The authors declare no competing interests.

## Supplemental Figures

**Figure S1.**
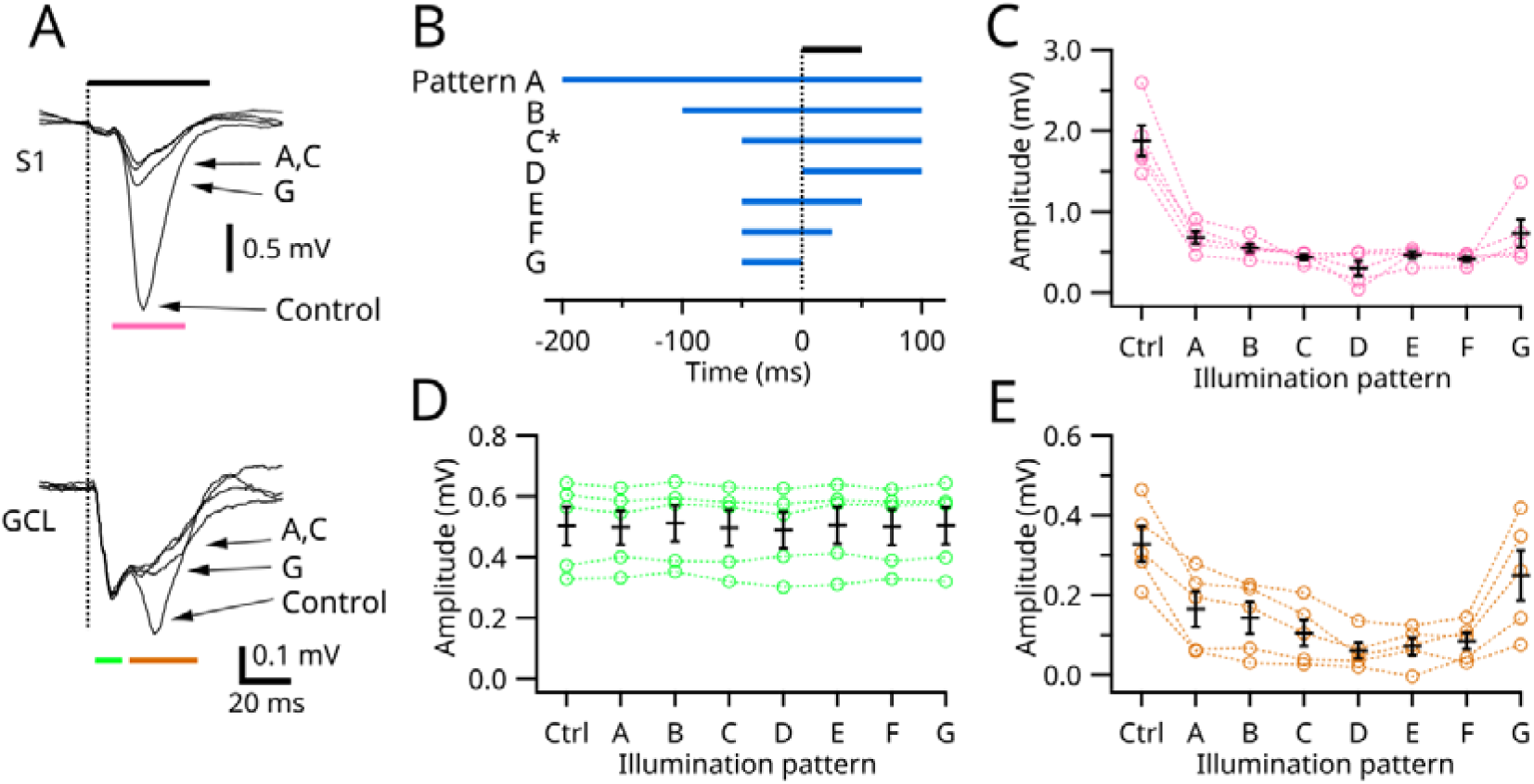
Exploration of temporal patterns of S1 photoinhibition. (A) Representative recordings of field potential in S1 (top) and cerebellar GCL (bottom). The averages from 20 traces in each stimulation pattern are shown. (B) Temporal patterns of the light illumination. The patterns B, D, E, and F are omitted in (A) for clarity. Black bar indicates the duration of air puff, whose onset is indicated by the vertical dotted line. (C–E) The peak amplitudes of S1 field potentials (C) and the early (D) and the late (E) components of cerebellar GCL field potentials (*n* = 5). Means ± SEMs are presented as black bars and lines, respectively.

**Figure S2.**
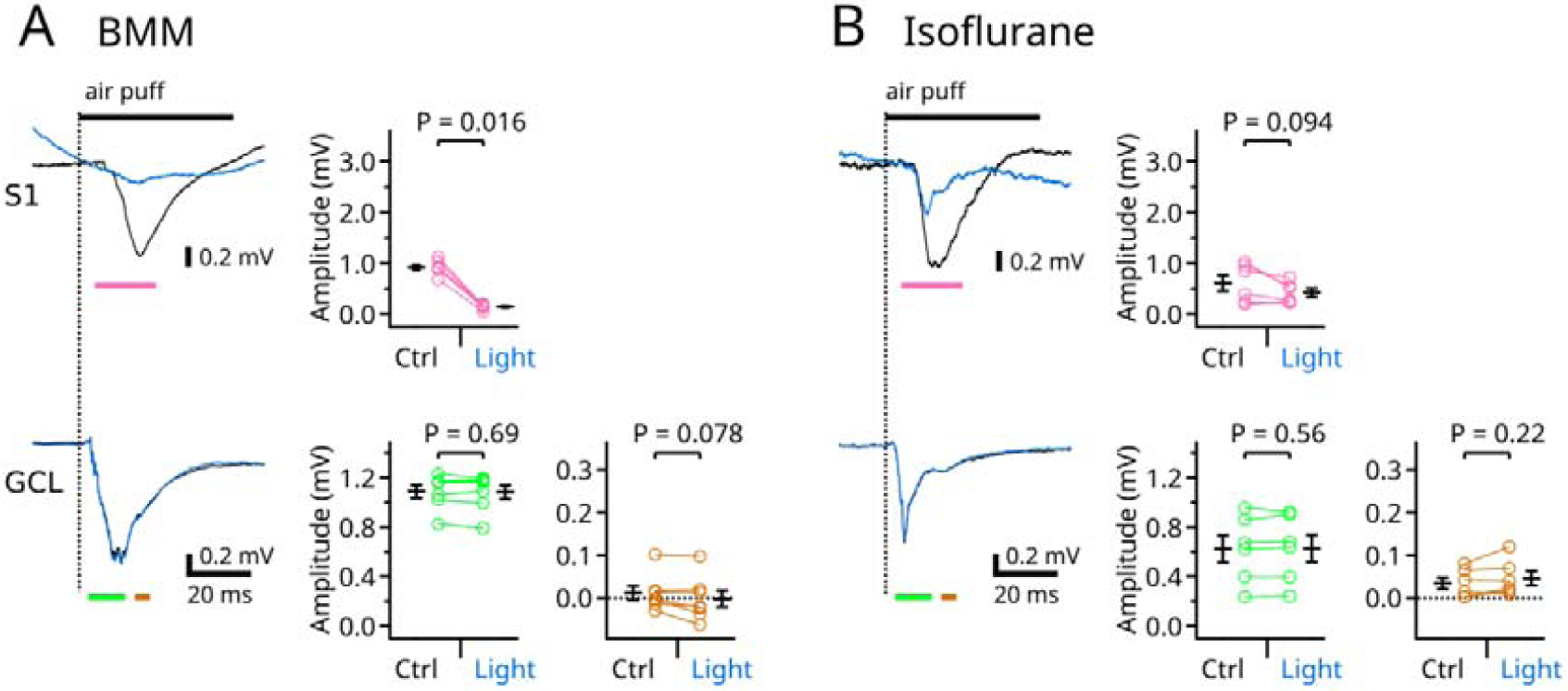
Effects of anesthetics on the cerebral and cerebellar responses. (A) The same experiments as described for Figures 1E and 1F, but the mice were anesthetized with a mixture of medetomidine (0.3 mg/kg), midazolam (4 mg/kg), and butorphanol (5 mg/kg) (BMM). (B) The same as in (A), but the mice are anesthetized with isoflurane (2%). Means ± SEMs are presented as black bars and lines, respectively.

**Figure S3.**
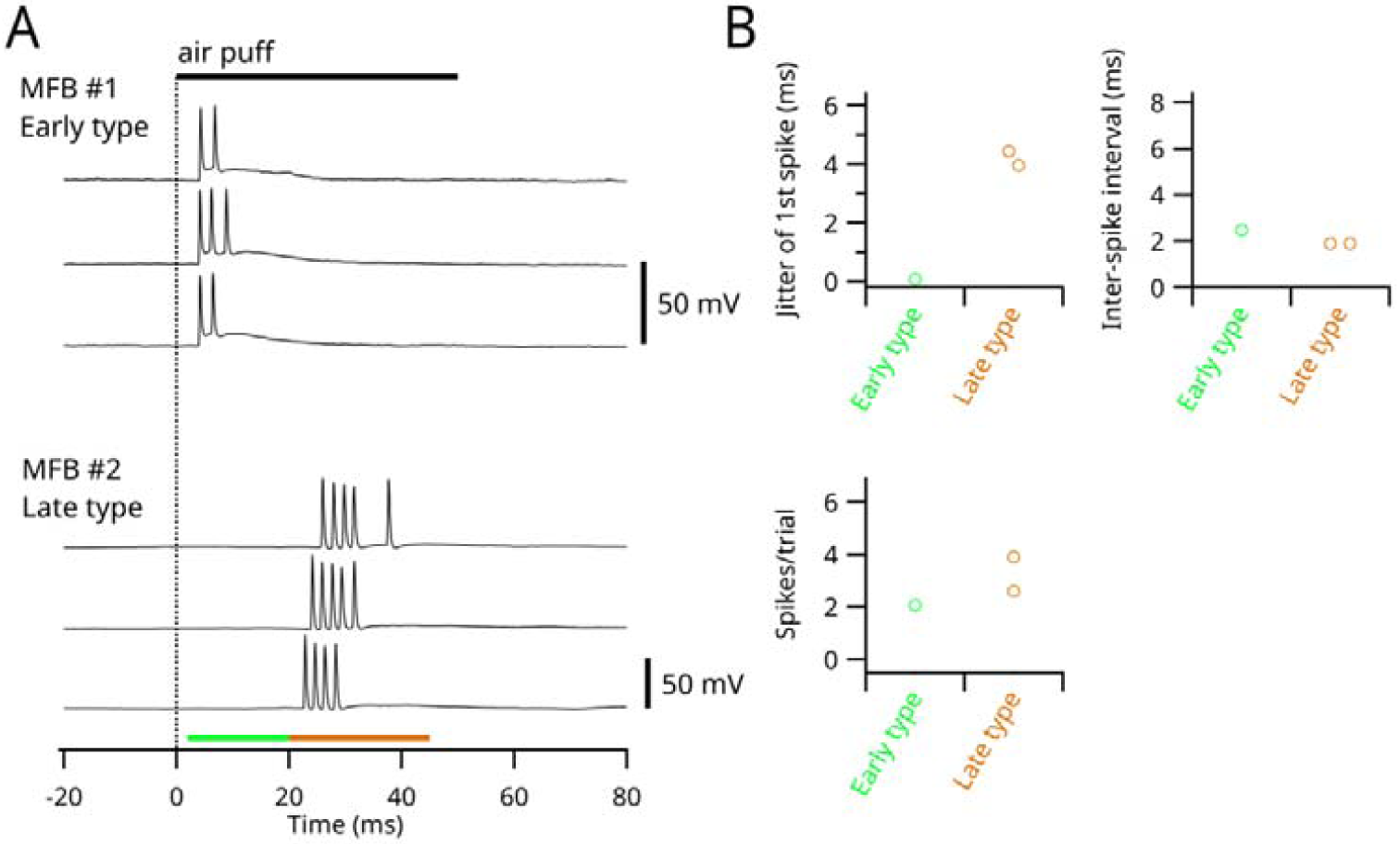
Sensory responses in putative mossy fiber boutons. (A) Whole-cell current clamp recordings from putative mossy fiber boutons (MFB; *n* = 3). MFB #1 was from a VGAT-ChR2 mouse, whereas MFB #2 was from an Aldoc-Venus mouse. Firing of MFB #1 was not affected by photoinhibition of S1 (not displayed). Three sample traces from each MFB are shown. MFBs were categorized as the early or the late type on the basis of their timing of firing. (B) Spike patterns were analyzed in the same manner as described for Figures 2D, 2E, and 2F.

**Figure S4.**
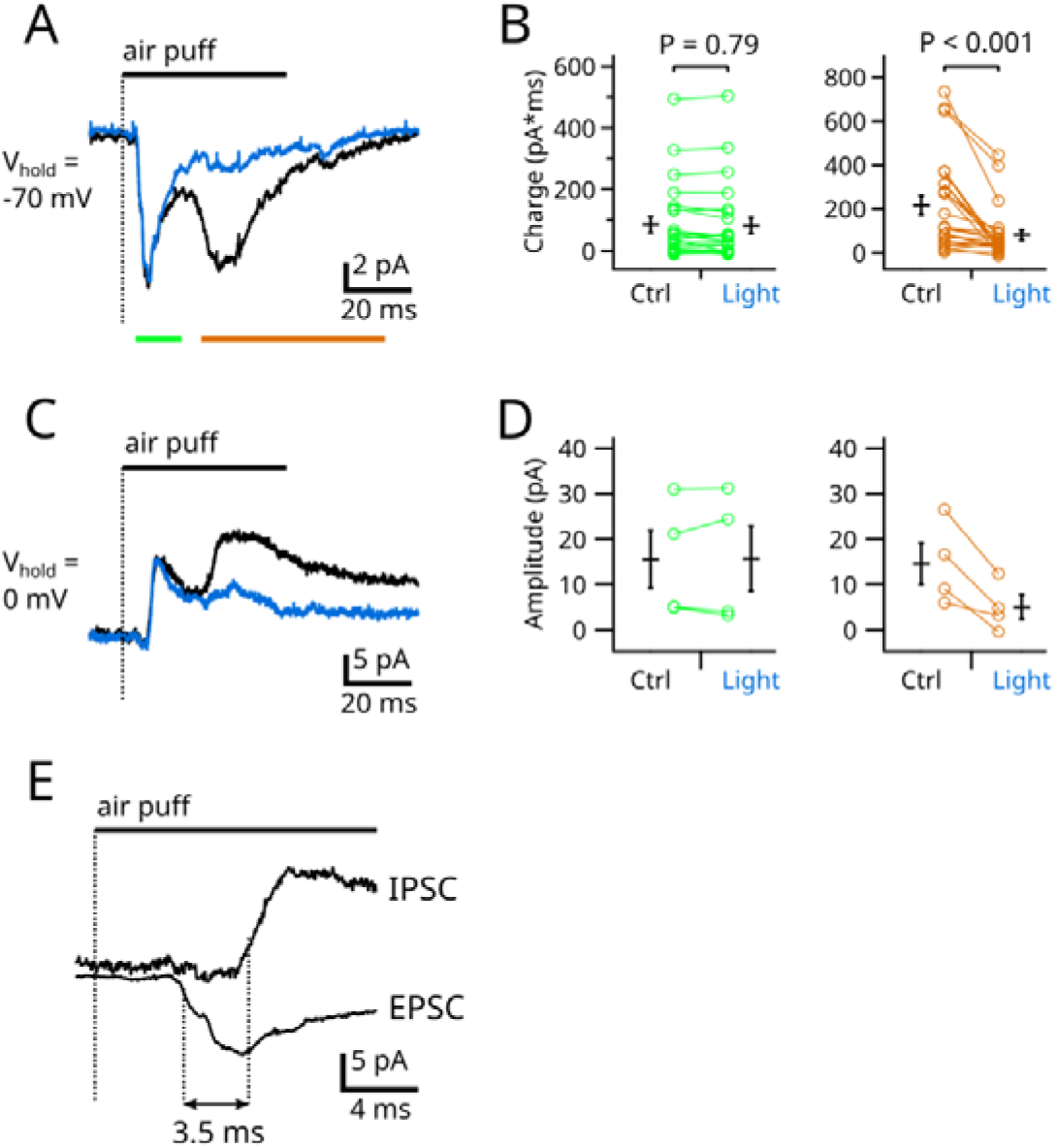
Extra data from whole-cell voltage clamp experiments. (A) Average traces of EPSCs in all 24 granule cells presented in Figure 2. (B) Synaptic charges (integral of the synaptic current, *n* = 24) measured for the time periods indicated in (A). The synaptic charges during the late phase (right, brown) may contain the tails of the early components (left, green). (C) Average traces of IPSCs from four granule cells voltage clamped at 0 mV (the reversal potential for the glutamatergic current). (D) The amplitudes of IPSCs (*n* = 4). (E) Comparison of onsets of an EPSC and IPSC. Control traces in (A) and (C) are shown together. The onset (20% rising point) of the averaged traces had a 3.5 ms difference. Means ± SEMs are presented as black bars and lines, respectively.

**Figure S5.**
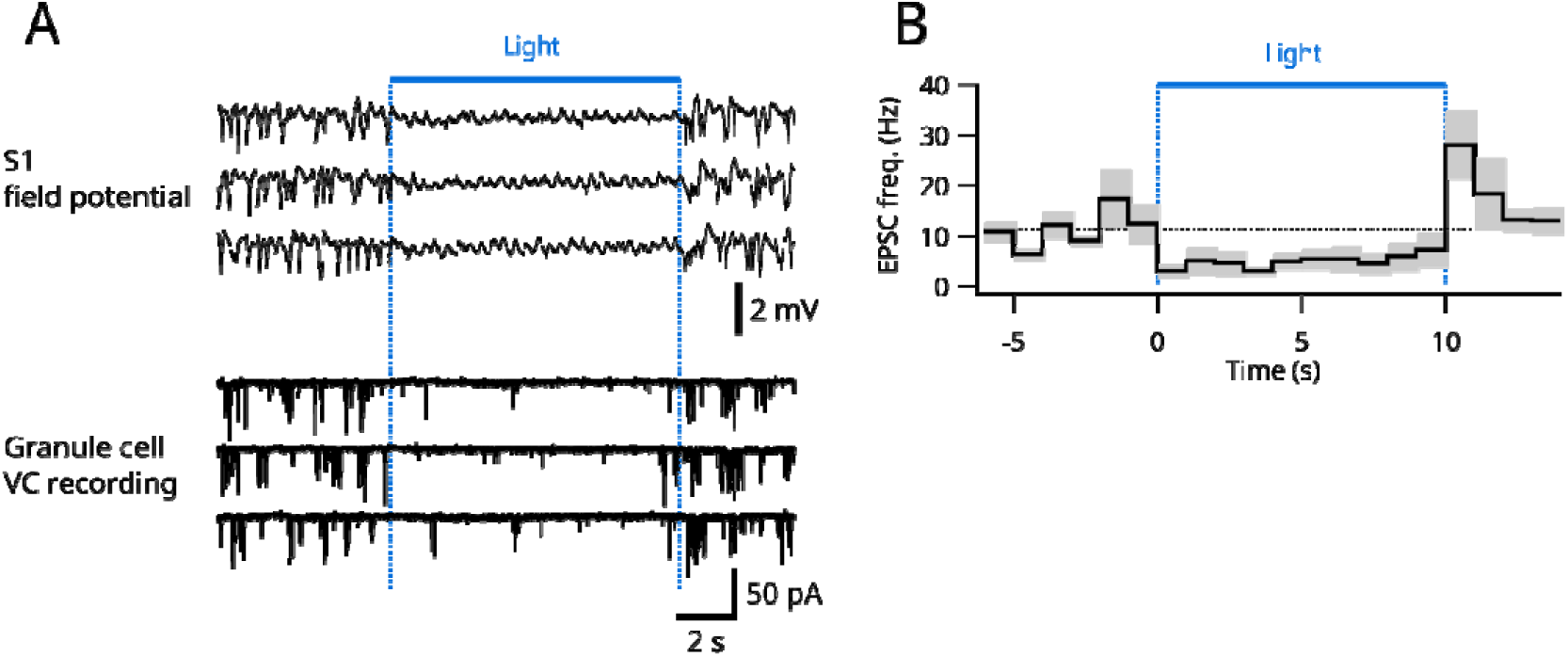
Effect of S1 photoinhibition on spontaneous EPSC in granule cells. (A) Representative recordings; simultaneous field potential recordings from S1 (top) and whole-cell voltage clamp recording from a granule cell (bottom). Long (10 s) blue light applications were repeated five times, but three trials are displayed. (B) Frequencies of spontaneous EPSCs (average from three granule cells) are displayed. Shading indicates SEMs.

**Figure S6.**
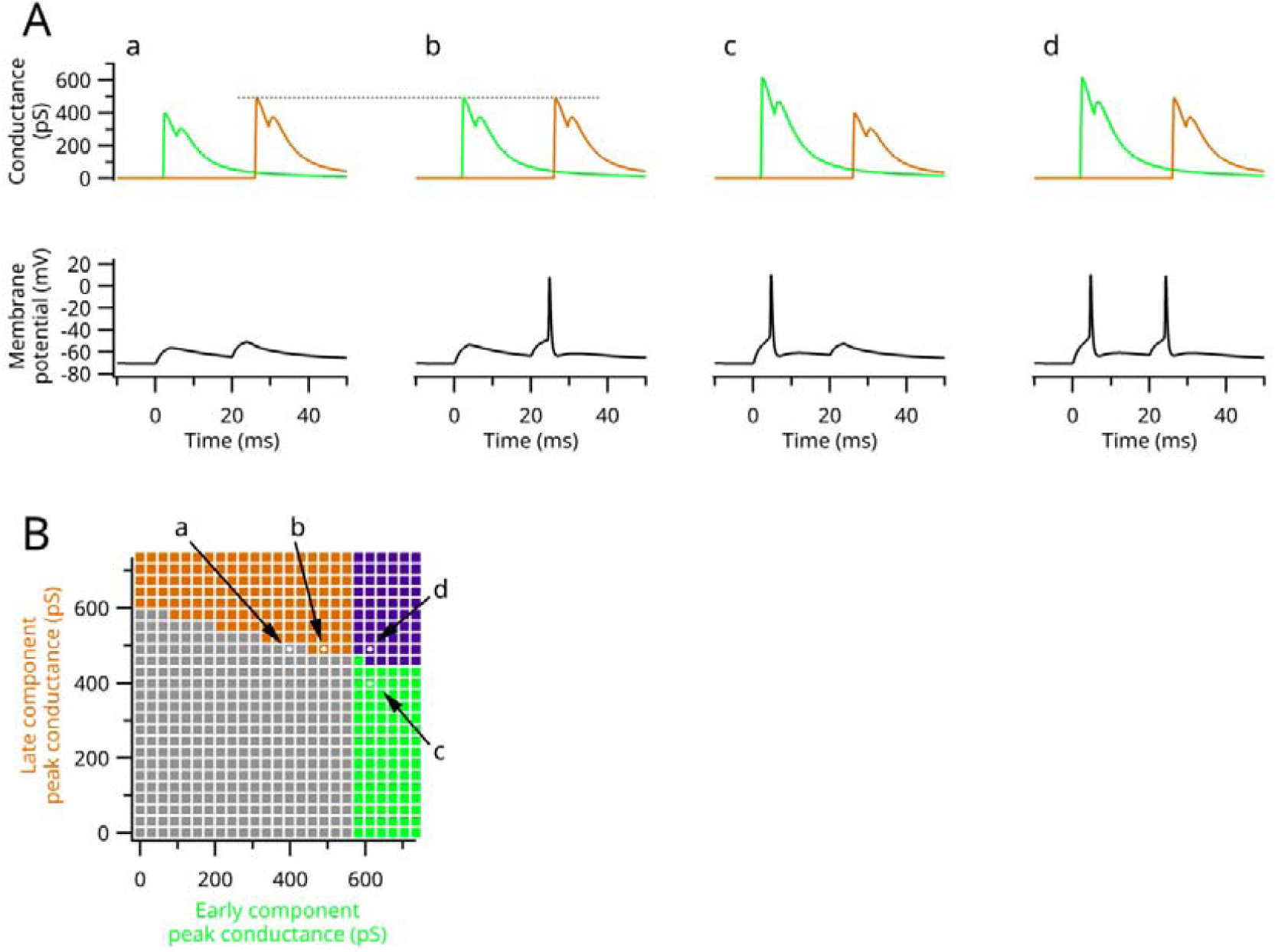
Computer simulation of action potential generation in granule cells. (A) Four representative patterns of simulation. Top, AMPA-type synaptic conductance. Bottom, resulting membrane potential at the soma. A realistic model of a granule cell was used as published by (Diwakar et al., 2009) for the NEURON simulation environment (Hines and Carnevale, 1997) without modifications except for AMPA and NMDA conductances, which were systematically changed while the NMDA:AMPA ratio was kept constant (0.2) (Cathala et al., 2003; Powell et al., 2015). Excitatory synaptic conductances, each of which consists of two synaptic events with a 3 ms interval, were applied to two dendrites with a separation of 20 ms. Inhibitory synaptic conductances, each of which consists of a single synaptic event with a 3.5 ms delay from the onset of excitatory synaptic conductance, were applied with a separation of 20 ms and a constant maximal conductance (756.35 pS). (B) The overall profile of spiking patterns triggered by various synaptic conductances. A plot resembling that in Figure 2B, but where the colors indicate the resulting spike patterns. White dots indicate the cases presented in (A).

**Figure S7.**
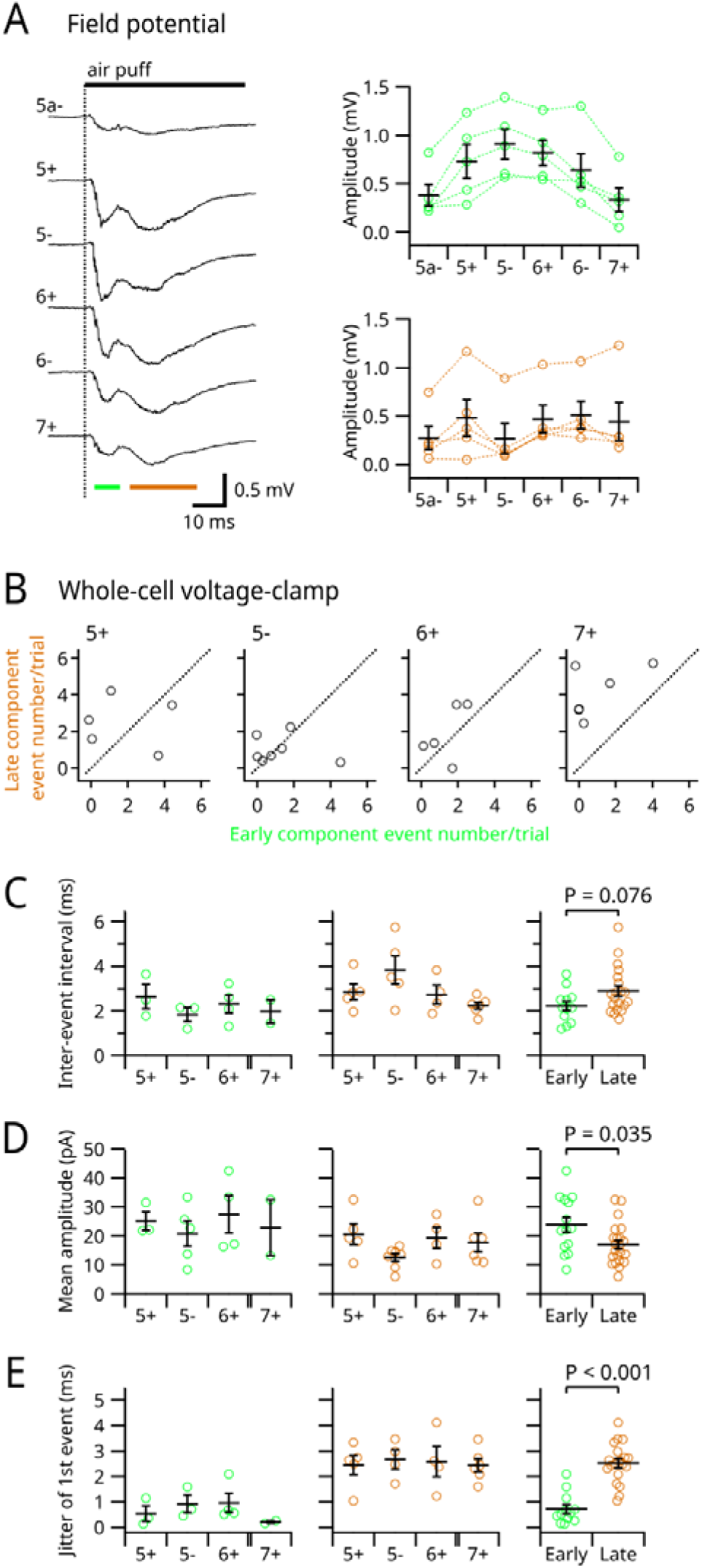
Extra data from experiments using Aldoc-Venus mice. (A) Left, representative field potential recordings from six different bands in crus II. Right, pooled data from five mice. The amplitudes of early (top) and late responses were measured during the times indicated by the colored bars in the left panel. (B) The numbers of evoked EPSC events in the early and the late phases were compared in individual bands. The same data as in Figure 4C were used. Dotted lines indicate unity. (C–E) The median interevent intervals (C), the mean amplitudes (D), and the jitter (the standard deviation of the timing) (E) of EPSCs in various bands in the early (left) and the late (right) phases. The right panels show pooled data from all bands. Means ± SEMs are presented as black bars and lines, respectively.

**Figure S8.**
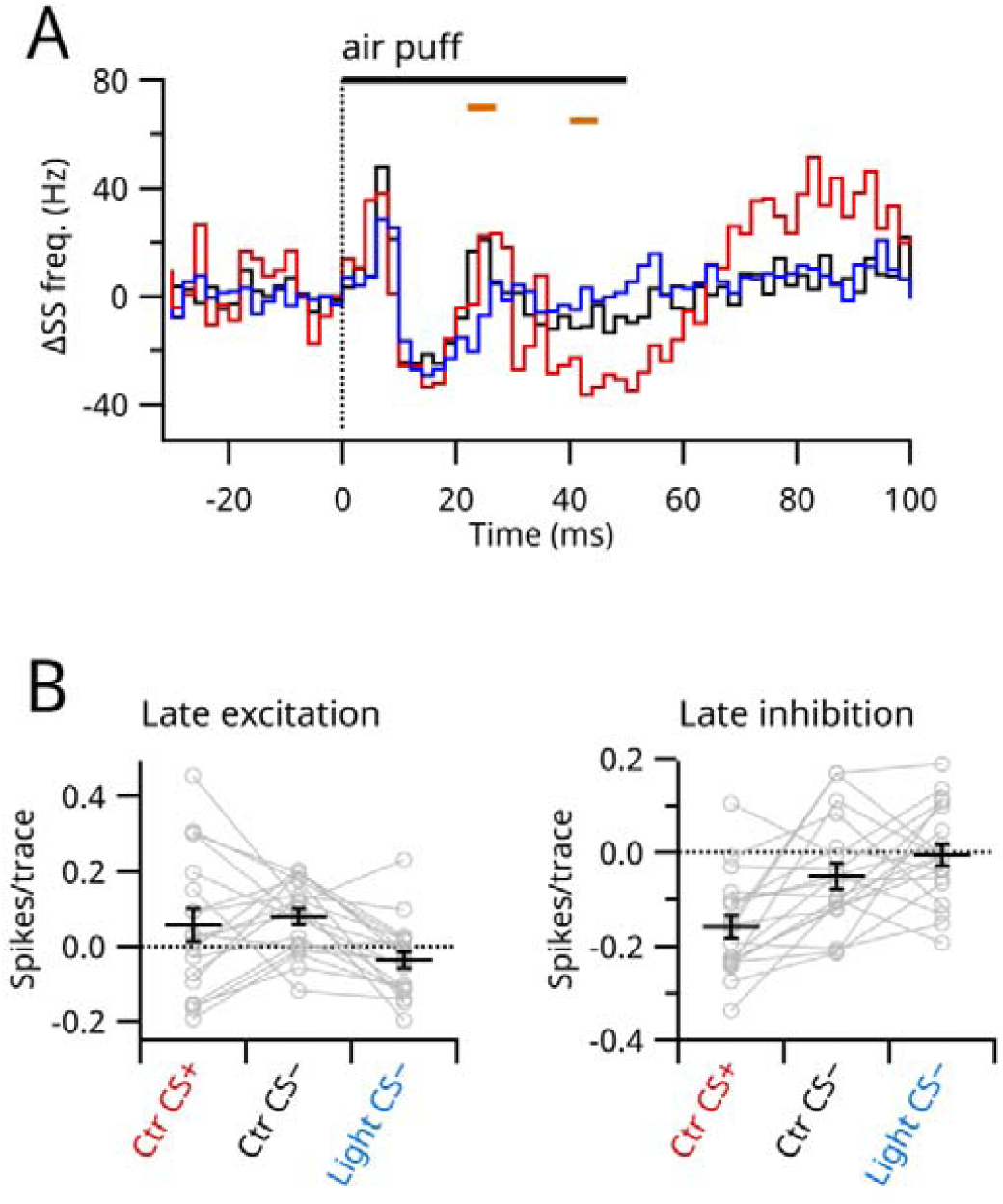
Detailed analysis of the interaction of SS and CS in Purkinje cells. (A) Average time histogram (2 ms bin, *n* = 19) of SS from trials with CS responses (red) and without CS responses (black) under the control condition and those without CS responses during photoinhibition of S1 (blue) are shown. In S1 photoinhibition, trials with CS responses were rare and therefore omitted in this analysis. (B) SS numbers were measured during the times indicated by bars in (A), the same timings as in Figure 5C. Means ± SEMs are presented as black bars and lines, respectively.

**Figure S9.**
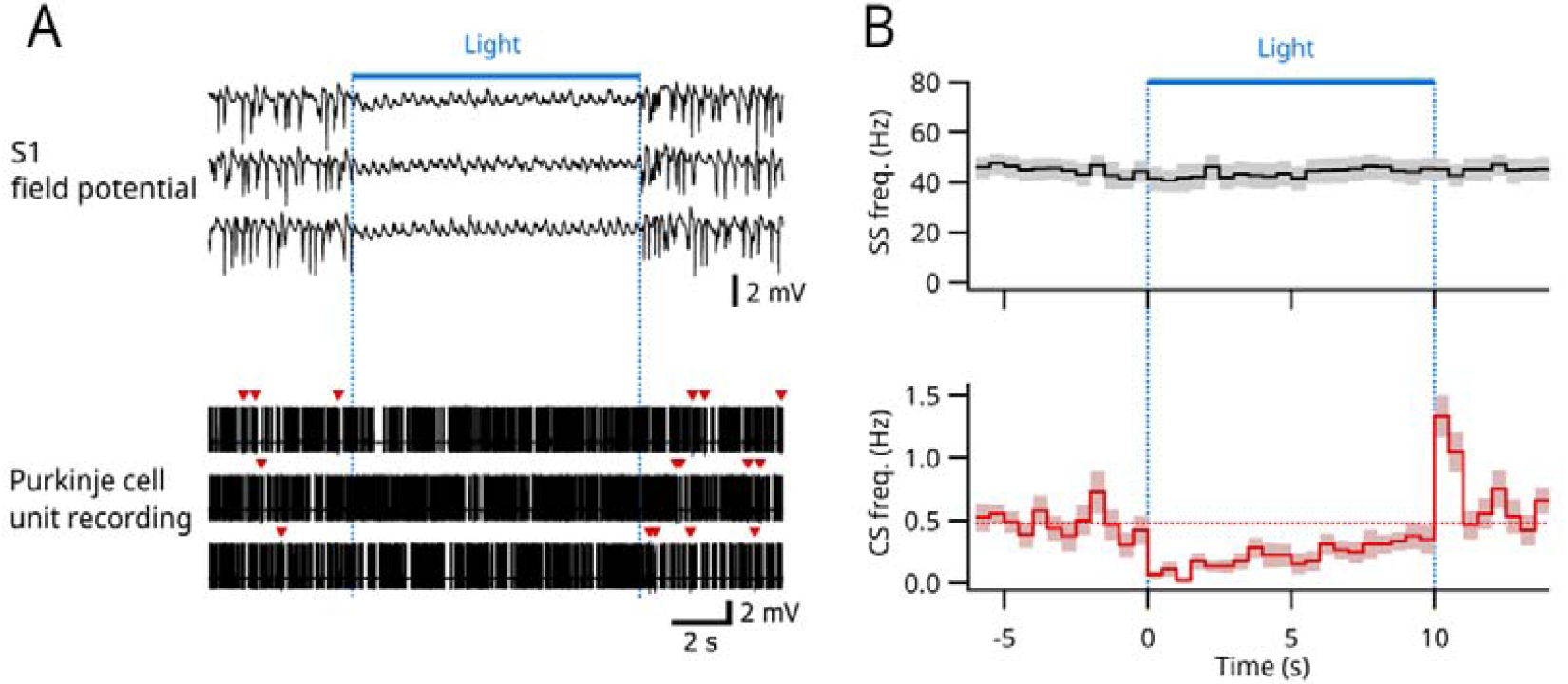
Effects of S1 photoinhibition on spontaneous SS and CS in Purkinje cells. (A) Representative recordings; simultaneous field potential recordings from S1 (top) and unit recordings from a Purkinje cell (bottom). Long (10 s) blue light applications were repeated six times, but three trials are displayed. Red arrowheads indicate CS. (B) Averages from 18 Purkinje cells. Spontaneous frequencies of SS (top) and CS (bottom) are displayed. Shading indicates SEMs.

